# Physiological predictors of reproductive performance in the European Starling (*Sturnus vulgaris*): II. Multivariate Analysis

**DOI:** 10.1101/330290

**Authors:** Alan A. Cohen, Mélissa Paquet, Melinda A. Fowler, Véronique Legault, Tony D. Williams

## Abstract

Physiological variation is generally thought or supposed to underlie variation in fitness related traits in wild animals, including reproductive effort, reproductive success, and survival. However, physiological markers of individual quality have proven elusive. In this paper and its companion, we use data on 14 physiological parameters measured in 152 observations of 93 individual European starlings (*Sturnus vulgaris*) over 2 years in an attempt to understand how physiological variation relates to variation in current breeding productivity, future fecundity, and survival. The companion paper, focusing on univariate analysis, showed that individual physiological parameters have little relationship with these performance measures. Here, we used more sophisticated statistical approaches in an attempt to extract a multivariate signal from the biomarkers – physiological dysregulation as calculated via statistical distance, and a number of principal components analysis approaches. Broadly speaking, there was a surprising lack of association between physiology and performance: while some physiological summary measures were associated with some performance measures, the associations were not particularly strong or robust given the large number of statistical tests conducted. This implies either that there are relatively few links between physiology and performance, or, more likely, that the complexity of these relationships exceeds our ability to measure and model it, even using state-of-the-art statistical approaches. This is likely particularly true because our population was quite heterogeneous; we nonetheless urge caution regarding the over-interpretation of isolated significant findings in the literature.

## Introduction

Much of the field of physiological ecology is based on the premise that fitness, fitness components, and key life history traits have coherent physiological underpinnings, and that we can thus understand how selection could act on life histories by understanding the mediating role of physiology [1, 2]. Indeed, there are a number of clear examples of exactly this kind of relationship, particularly with regard to hormonal control of reproduction, reproductive behaviour, and risk-taking [e.g. 3, 4]. However, the same clarity has not been evident in studies looking for physiological markers of individual quality or condition. For example, in animal physiology, research into immune function and oxidative balance has grown quickly over the last 15 years, often with the objective of identifying individuals with more robust immune responses or lower levels of oxidative stress as correlates of other ecological or individual variables [e.g. 5, 6]. In both these fields, results have been mixed and confusing [7-13], particularly considering publication bias and other biases leading to hypothesis confirmation [14].

These conflicting results are starting to be understood from two angles, theoretical and empirical. From a theoretical perspective, ecologists have been naïve physiologists, simplifying complex, non-linear physiological systems into concepts (e.g. “oxidative stress,” “immunocompetence”) that may or may not have some basis in reality [15]. For example, oxidative stress is a combination of pro- and antioxidant forces [16], and each of these is a combination of many individual molecules that interact in complex feedback loops. Damage levels – and the importance of damage for organismal fitness – can vary greatly across tissues, cell types, cell compartments, membranes versus cytosol versus plasma, etc. Oxidative stress may be hormetic, inducing compensatory responses that benefit the organism under certain conditions [17, 18]. Despite knowledge of this complexity, research on oxidative stress in an ecological context usually uses one or two markers under the supposition that these can tell us most or all of what we need to know [19-21].

Complementing this from an empirical perspective is the consistent finding that correlations among purported markers of a given over-arching concept (e.g. oxidative stress) are highly unstable over time, populations, species, conditions, analysis levels, etc. [13, 22, 23]. How are we to understand “antioxidant capacity” when at least three independent axes are needed to summarize the variation in circulating micronutrient antioxidant levels (even without considering enzymes or tissues), and when the composition of these axes is unstable [24]? This problem is reflected in highly contradictory results observed throughout the literature.

In response to this problem, a number of approaches have been proposed, mostly focusing on multivariate analysis of physiological markers to obtain a more stable, clear signal of underlying processes. Principal components analysis (PCA) is the obvious first option, though as mentioned it tends to identify, but not solve, the problem of multi-dimensional variation and instability of axes [12, 13, 23, 24]. More recently, we proposed an approach based on statistical distance to measure body condition as a function of how close or far an individual is from a homeostatic body state norm [25-27]. Briefly, using Mahalanobis distance (DM), an individual has a score of 0 (optimal condition) if the individual has the mean levels of all biomarkers; scores increase as distance from this centroid goes up in multivariate biomarker space (the individual has a more “abnormal” profile). DM automatically adjusts for the correlation structure among biomarkers, down-weighting distances along major axes of correlation and up-weighting those against these axes. This is crucial because the correlation structure of the biomarkers represents the most normal ways the markers co-vary. For example, if height and weight were considered independently, we would falsely conclude that it is equally likely to be tall and heavy as short and heavy, but considering the correlation we can accurately identify the short-heavy combination as the rarer one. While in theory it might seem that mean biomarker levels are a poor proxy for ideal levels, in practice this seems to pose minimal problems when enough markers (10-15) are included [25]. DM has been extensively validated as a measure of physiological dysregulation during human aging [28-32], and was also predictive of body condition in a wild-caught population of a shorebird, the red knot (*Calidris canutus*) [27]. However, it has not yet been validated in other ecological applications.

Use of individual markers may also have advantages: they may be more intuitive for readers, easier to interpret biologically, and more specific about identifying mechanisms. In the companion paper [33], we examined levels of 13 individual biomarkers in relation to reproductive effort (workload) and breeding performance in a population of European starlings (*Sturnus vulgaris*) followed over 2 years, finding essentially no association between levels of the individual biomarkers and levels of reproductive effort or performance. Physiology did, however, vary across year and breeding stage. This is an example of the negative results that are often harder to publish but (anecdotally among colleagues) are very common in this field. In our case, this is despite a relatively large sample size (76 individuals, 152 data points), a diversity of physiological measures, and a broad array of effort and reproductive performance measures (current productivity, future fecundity, and survival). Here, we ask whether our negative finding with the individual markers could be attributable to a lack of appropriate multivariate analyses integrating them. It was not apparent *a priori* whether such analyses were appropriate or necessary: we could plausibly have found that one biomarker was highly predictive and that others diluted the signal. We generate integrated measures of physiological condition using both PCA and DM implemented under a variety of suppositions. We also perform multivariate analysis (largely PCA) on the reproduction variables in an attempt to extract a stronger signal. We predicted that high DM would be associated with low reproductive success. Because of the complexity of both the data structure and the analyses, we present detailed methods and results in the S1 Appendix, with an overview of typical findings presented in the main text.

## Materials and Methods/Results

### Overview

Because our analyses are highly exploratory and complex, many methodological decisions depended on intermediate results. We thus take the unusual approach of presenting a joint Methods-Results section, so that the logic of our methodological decisions can be seen in light of preceding results. Despite the exploratory nature of our analyses, we had a clear general *a priori* plan, as outlined by the following steps:

1) Process individual physiological variables via transformations, outlier detection, and PCAs (if justified) to combine markers within systems.
2) Generate a simple version of DM based on the population mean
3) Generate more sophisticated versions of DM based on *a priori* knowledge of biomarkers
4) Generate PCA-based scores to summarize overall variation in physiology.
5) Generate a series of composite variables that summarize variation in reproductive success.
6) Test for associations between each composite physiological variable and the raw and composite performance variables
7) Interpret the overall results in the light of general patterns across the analyses, not the significance of individual tests. For example, do some versions of DM show a consistently stronger signal? Are associations with DM consistent between raw and composite measures of reproductive success, and across versions of DM? Are associations detected particularly strong and/or more frequent than expected based on false discovery rates? Multiple testing is thus not formally adjusted for, but our approach is conservative overall.

Note that statistical analyses in this paper and the companion paper were conducted by different team members, and while we have discussed multiple analytical decisions, we have not tried to perfectly harmonize results. We believe that many analytical decisions (log transformation, outliers, etc.) have no single correct response, and that it is thus more scientifically honest to allow for modest variation between the papers. Major discrepancies were adjudicated. All analyses were conducted in R v. 3.0.0 and 3.2.2.

### Data

The study population of European starlings is described in substantial detail in the Supplementary Material. Here we highlight only the key features relevant to this analysis. The overall structure of the study is shown in Fig. 1. Breeding females were followed in 2013 and 2014. Blood samples were taken during incubation, chick-rearing, and (when relevant) chick-rearing of a second brood. Blood samples were analyzed for 14 physiological variables, two of which were combined into a single index in the companion paper. Current productivity was measured via brood size at day 6 and at fledging (day 21) and chick mass measured at day 17 for each brood, as well as composite measures. Future fecundity was measured via the same variables in 2014, for individuals present in both years. Survival was measured as local return rate. Details of data collection are provided in the S1 Appendix. All research was conducted under Simon Fraser University Animal Care permits # 657B-96, 829B-96, 1018B-96).

**Figure 1.**
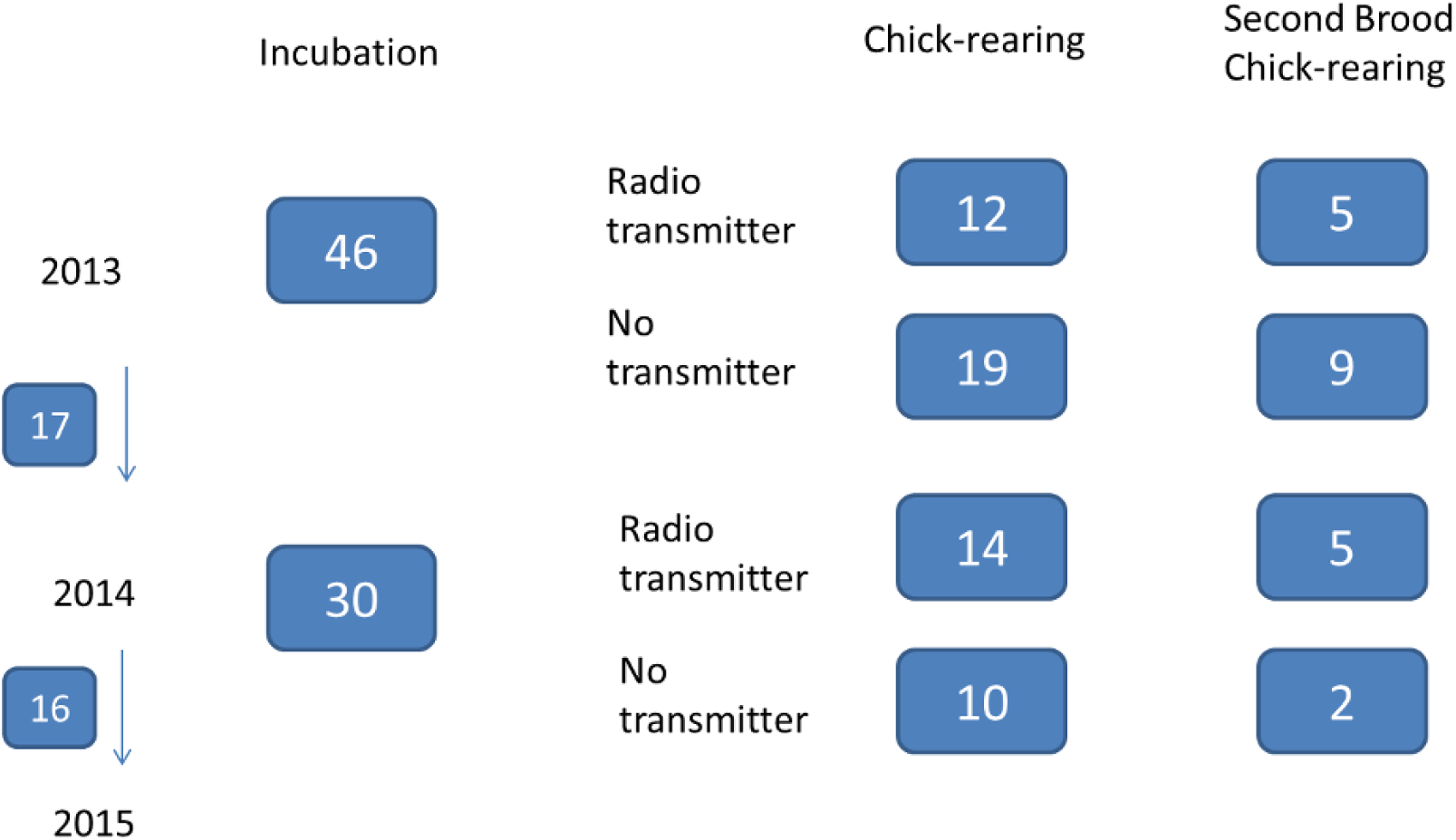
Experimental design.

Overall, the sample size (152 data points) is relatively large for a dataset with this depth of both physiological and reproductive data. However, that number falls substantially when considering that complete physiological data are needed for many of our analyses; only 80 data points have all 14 measures, though 24 more are available if the reticulocyte variable is excluded. As can be seen in Fig. 1, the sample size is large if subgroup analyses can be ignored; however, many subgroups are too small to permit independent analyses. We thus pooled first and second brood chick-rearing data into a single “chick-rearing” category. We chose to include data from both first and second broods to increase our sample size and to ensure that we fully captured the breeding success of the most successful individuals, but are aware that it might complexify the interpretation. Nevertheless, we ran analyses excluding second broods and obtained highly consistent results (data not shown).

### Pre-treatment of physiological variables

All physiological variables were assessed for normality and log- or square-root transformed as necessary, and we tested extensively for outliers (see S1 Appendix for details). We looked at possible effects of from when box was plugged early in the morning to when bird was removed on physiological variables and found no clear evidence for such effect (see S1 Appendix *section 3.1* and S1-S2 Figs.). In order to understand the variation in oxidative stress and immune function and potentially reduce the number of variables, we performed separate 2-variable PCA analysis on the antioxidant level (OXY) and dROMs measures (oxidative stress) and the agglutination (AG) and lysis variables (immune function). In both cases, we detected a similar and surprising structure: the first axis loaded in the same direction with both variables (78% and 62% of variance explained, respectively; see S3 Fig.), whereas the second axis explained the contrast between the variables (one negative and one positive loading). This was particularly surprising for OXY/dROMs, where we expected the first axis to explain the contrast, with high OXY and low dROMs indicating “good” oxidative status; instead, our axes appear to indicate that OXY tracks dROMs, with high on both presumably indicating a lack of control of dROMs and concomitant up-regulation of OXY (S3 Fig., panel b). For both the immune and oxidative stress variables, we used the two PCA axes rather than the original variables because the biological interpretation is clearer: PCA1 is activation of the system (higher levels of both markers) and PCA2 is balance of the system (relative levels). In both cases, we also tested ratios of the raw variables and found they were strongly associated with PCA2, confirming this interpretation (S3 Fig.).

Because of the possibility that physiology varied systematically across year and breeding stage, we tested this for each of the 14 physiological markers using t-tests. (Similar tests were conducted using mixed effects models in the companion paper.) Overall, we obtained results very similar to the companion paper [33] showing major changes in physiological parameters across both year and breeding stage, though there were significant year-stage interactions in only three parameters. Accordingly, we decided to perform our standardization by year and breeding stage. This makes statistical sense, and could have a conceptual justification as well, if optimal physiological state depends on factors that change substantially across years or breeding stages. However, sample sizes for the four year-stage subgroups were too small to produce robust estimates of means for standardization (S4-S5 Figs.). We thus used regression to identify beta-coefficients with which to adjust year and stage means. Biomarker values used in all subsequent analyses thus reflect the number of standard deviations away from the mean, given the year and stage at sampling.

### Basic construction of DM

DM was calculated using the following formula:

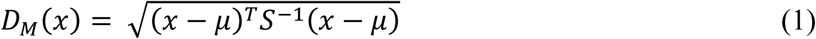

where *x* is a multivariate observation (a vector of simultaneously observed values for the variables in question, such as all the biomarker values for a given bird at a given time point), *μ* is the equivalent-length vector of sample means for each variable, and *S* is the sample variance-covariance matrix for the variables. Because the sample in this case is other birds in the same year and stage, we used the regression-based adjustments noted above to calculate *μ* specific to each year and stage. We used a single global correlation matrix *S*, which appeared sufficiently robust that it did not require year- and stage-adjustments, and in fact would have suffered from the smaller sample size (S4-S5 Figs.). We tested the robustness of DM to removal of individual markers (S6 Fig.) and found that it was highly correlated with the full version regardless, indicating minimal sensitivity to any particular marker included. We had hoped to exclude the reticulocyte variable due to the missing data noted earlier, but among biomarkers DM was slightly more sensitive to inclusion/exclusion of this variable; we thus present analyses with and without reticulocytes included, and this sometimes changes results.

### Centering DM

DM is defined as a distance from a centroid, normally the population mean. As applied to physiology, the centroid is supposed to represent the ideal physiological/homeostatic state. Previous studies have found the mean to be a reasonable approximation of this optimum for most biomarkers, and DM shows little sensitivity to the precise definition [25, 26]. However, in this case we had a number of biomarkers for which there was a clear *a priori* expectation that the optimal value was not the mean, in contrast to all previous studies. For example, low scores of the two OXY/dROMs PCs represent low activation of oxidative stress pathways and balance toward antioxidants rather than free radicals, respectively, so lower scores should be better, with no lower bound. For some other variables, we had a hypothesis about direction but no certainty. We thus used a combination of *a priori* knowledge and data driven methods to generate multiple versions of DM (approximately 20) based on which biomarkers were considered optimum at their mean, minimum, or maximum. However, DM version has relatively little impact on our overall conclusions (see section *Centroid calculation* in S1 Appendix for details), and we thus chose to present several of the most distinct, for ease of presentation. S1 Table summarizes the final set of DMs that we kept for further analyses, including the basic one with year- and stage-specific means as the centroid. S7 Figure shows correlations between these different versions (see S1 Appendix for details).

### Physiological PCA

We conducted PCA in two general ways: on all variables, and on subsets identified through *a priori* knowledge (e.g. S8 Fig.). For the PCA on all variables, we had two versions: (1) PCA on the 14 standard normal variables, and (2) PCA on the log of the absolute value of the 14 standard normal of each variable (i.e., looking for a structure in the deviance, related to the theory underlying DM; S9 Fig.). However, the first axes explained only 14% and 13% of the variance respectively, and 6/14 of the axes had eigenvalues greater than 1 in both cases, suggesting minimal efficacy of the PCA at reducing the dimensionality. Examination of loadings also did not provide a clear biological interpretation. Likely the combination of physiological complexity and small sample size given the number of variables made results unstable; we nonetheless tested associations between the first three axes of both PCA and the performance variables for exploratory purposes (see below). We have previously found that some physiological PCA axes with small variance explained and unclear biological interpretation can nonetheless be highly stable across populations and predictive of health status [34].

For the PCA based on *a priori* knowledge, we conducted three analyses. The first was intermediary metabolism and energy supply (“metabolism/energy”) and included glucose, NEFA, uric acid, and triglycerides (S10 Fig., panel a). The first axis explained 37% of the variance and contraposed glucose (+) against the other three (-). The second analysis was on aerobic/metabolic capacity (“aerobic capacity”) and used hematocrit, hemoglobin, reticulocytes, and cort (S10 Fig., panel c). The first axis explained 36% of the variance and contraposed cort against the other three. The third analysis was on oxidative stress and muscle damage (“oxidative stress”), and used OXY, dROMs, and creatine kinase (S10 Fig., panel d). The first axis explained 63% of the variance and strongly contraposed creatine kinase against OXY and dROMs. The first axis from each of these three analyses was used in subsequent analyses.

### Performance variables

Due to missing data and biological complexities of the data structure (e.g. many individuals did not have a 2^nd^ breeding attempt), we performed PCA on only 4 variables reflecting current breeding productivity: first brood size at day 6, first brood size at fledging (day 21), total young fledged in the year, and total mass of fledglings produced over the year. The first axis explains 58% of the variance and is strongly negatively associated with the 3 variables describing number of young, but not fledgling mass. The second axis explains 26% of the variance and is nearly entirely explained by mass, which loads at 0.978.

### Associations between integrative physiology and performance variables

We conducted separate analyses on incubation physiology in relation to performance variables, chick-rearing physiology in relation to performance variables, and mean physiology across incubation and chick-rearing in relation to performance variables. We predicted that high DM would be associated with low reproductive success. We ran linear mixed effects models using the lmer function in the lme4 package [35] with the performance variable as the dependent variable, the physiological measure as the independent variable, and individual as a random effect. lmer does not calculate p-values; we approximated p-values by averaging the p-values calculated by taking the degrees of freedom based on the number of individuals and the degrees of freedom based on the number of data points; in this context, the difference is minimal and precise p-values are not important.

Overall, DM appears to be largely unassociated with performance variables (Fig. 2). Only 23 tests of 396 in Fig. 2 (5.8%) were significant at α=0.05, not much more than expected by chance. Moreover, the tests are not independent (there are strong correlations among DM versions and among performance measure subsets, see above). During chick-rearing, four of six DM versions show a clear negative association with total young fledged in a year (BSF sum per year) when the analysis includes reticulocytes, but this is replicated in only one DM version after excluding reticulocytes (Fig. 2), and is not replicated for incubation values. Further, this finding is not broadly confirmed by other reproduction variables, including the reproduction PC axis, fledgling mass, intermediate brood sizes, or subsequent year lay date. Given the number of tests we have performed, the underwhelming consistency, and the underwhelming p-values, we cannot draw a clear conclusion for an association.

**Figure 2.**
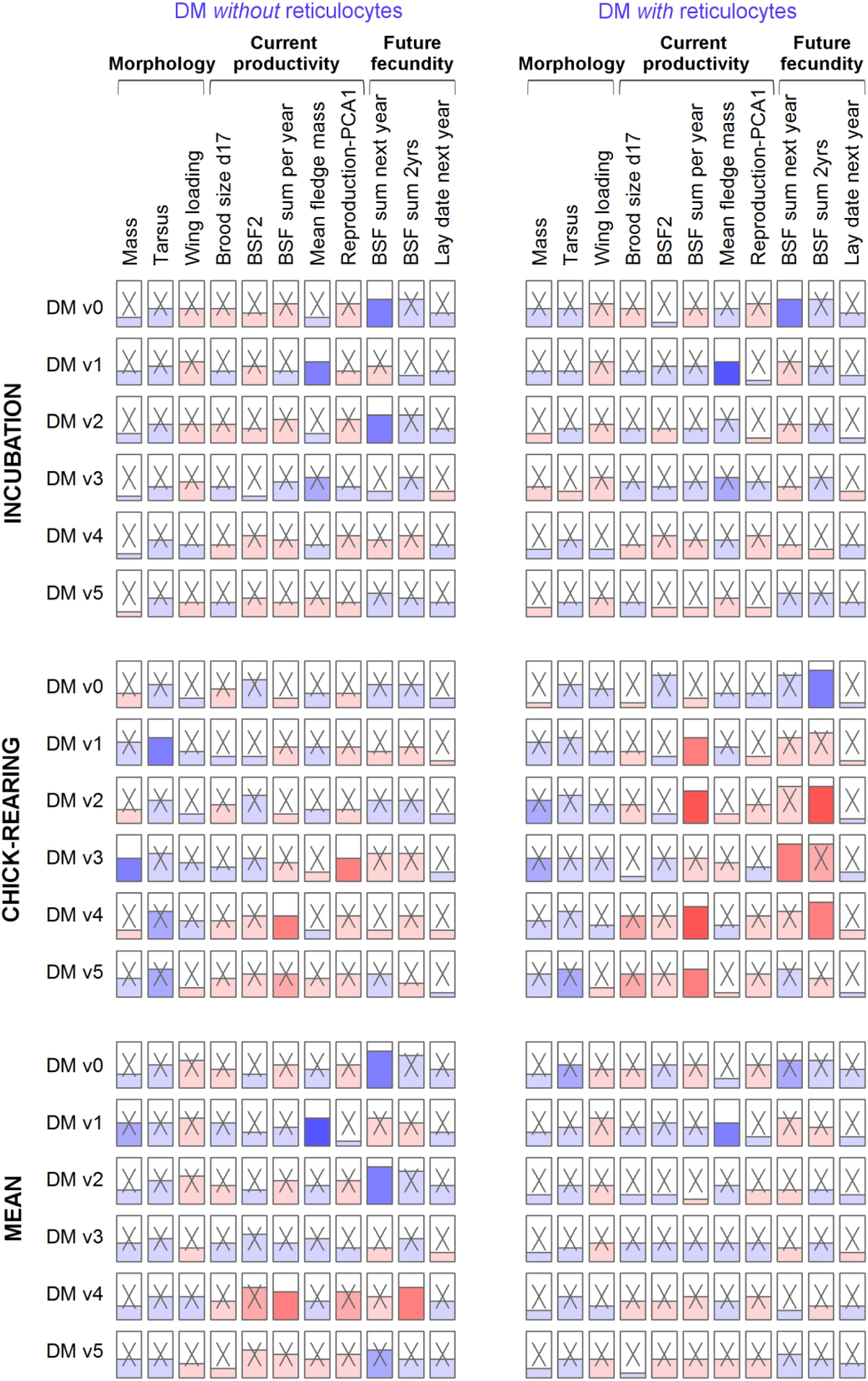
Few associations between physiological summary variables (DM) and reproductive performance. We performed regression models to evaluate the effect of different DM versions, excluding reticulocytes or not (n=104 and 80 respectively; see section *Centroid calculation* in S1 Appendix for details), on each performance measure, using observations during the incubation stage (**upper panel**), the chick-rearing stage (**middle panel**), or the mean of all available observations within a year (**lower panel**). Height of the horizontal line in the box represents the effect size and the color underneath is blue for positive effects and red for negative ones, with color hue illustrative of *p*-values (darker shades for lowest *p*-values). Non-significant coefficients are marked with an “X”.

To assess the association between DM and subsequent survival, we performed logistic regressions with the local return next year variable (glm function), which was not associated with any version of DM (Table 1; logistic regression results could not be included in Fig. 2).

**Table 1.** Survival analysis through logistic regression with local return next year, for six versions of DM (V0-V5).

| **Survival** |  |  |  |  |
| --- | --- | --- | --- | --- |
| **DM (n=35)** | **OR** | **LCI** | **UCI** | ***p*** |
| **V0** | 0.917 | 0.163 | 1.671 | 0.822 |
| **V1** | 1.126 | 0.384 | 1.869 | 0.753 |
| **V2** | 0.812 | -0.041 | 1.665 | 0.633 |
| **V3** | 0.744 | -0.124 | 1.613 | 0.505 |
| **V4** | 0.939 | 0.189 | 1.689 | 0.869 |
| **V5** | 0.816 | 0.112 | 1.521 | 0.572 |

Similarly, analyses relating physiological PCs to performance variables (Fig. 3) showed no robust associations. Some associations were significant, but there were no clear patterns and most or all of these are likely false positives due to the large number of tests performed (29 of 297 (9.8%) significant at α=0.05). Not a single positive result was replicated between incubation and chick-rearing analyses. Of course this does not exclude the possibility of some true positives among our significant results – PCA1 directional during incubation and PCA3 directional during chick-rearing seem to have disproportionate numbers of significant associations, for example – but the consistency of the findings is insufficient for us to draw firm conclusions.

**Figure 3.**
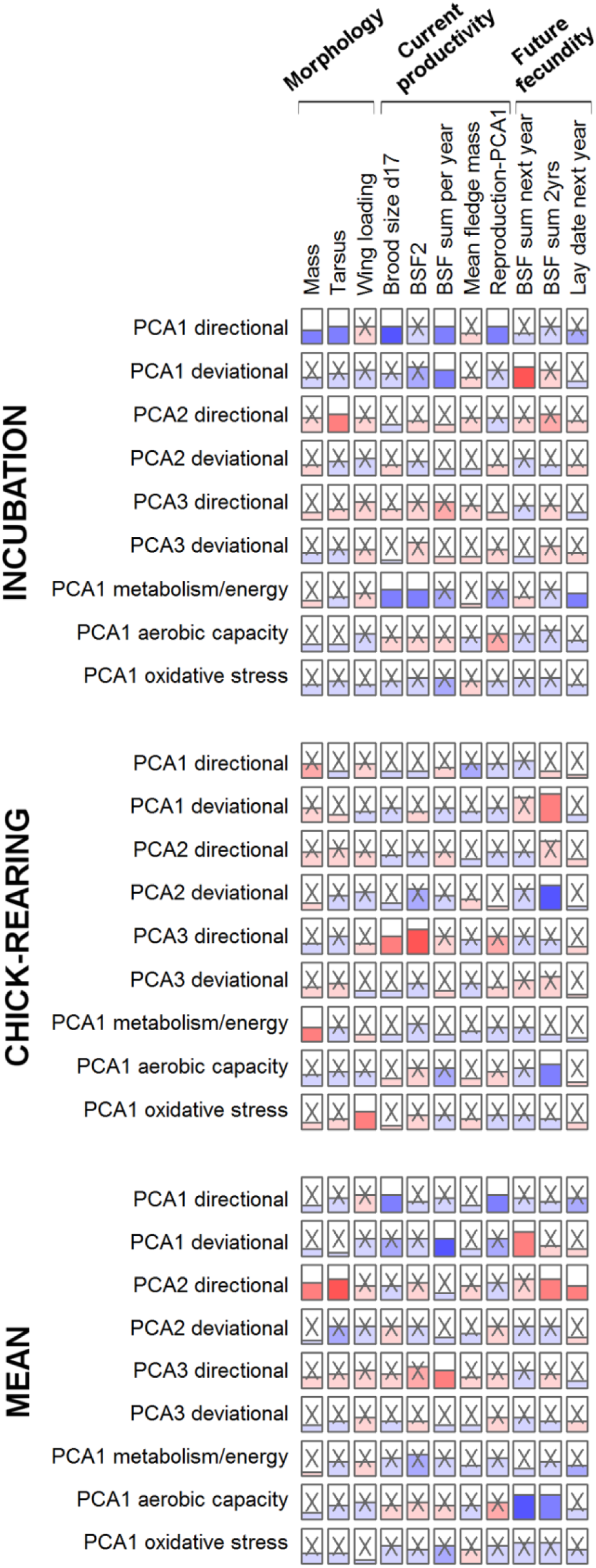
Few associations between physiological summary variables (PCA axes) and reproductive performance. We performed regression models to evaluate the effect of the first three axes of the PCAs on all centered and reduced physiological variables (PCA directional, n=80), on the logarithm of their absolute values (PCA deviational, n=80; see section *PCA with all physiological variables* in S1 Appendix and S9 Fig.), and the first axis of PCAs on functional groups of physiological variables (n=133, 92, and 122 respectively for metabolism/energy, aerobic capacity, and oxidative stress functional groups; see text and S10 Fig. for details), on each performance measure, using observations during the incubation stage (**upper panel**), the chick-rearing stage (**middle panel**), or the mean of all available observations within a year (**lower panel**). Height of the horizontal line in the box represents the effect size and the color underneath is blue for positive effects and red for negative ones, with color hue illustrative of *p*-values (darker shades for lowest *p*-values). Non-significant coefficients are marked with an “X”.

## Discussion

We constructed composite physiological variables using several multivariate methodologies and tested for associations between physiology and indicators of reproductive success. We were strikingly unable to find clear associations between physiology and individual quality in this dataset. While we cannot exclude the possibility of the predicted positive association between total young fledged and dysregulation scores (DM), the evidence is hardly convincing. The lack of clear associations is particularly surprising given the relatively large sample size for a data set of this kind and the relatively large and broad set of physiological measures. DM becomes more robust as more markers are used to calculate it [25]. These findings add to the negative finding of few or no associations between physiology and performance with the individual biomarkers in the companion paper (despite some systematic variation with year and breeding stage) [33].

It is of course possible that our negative results are due to a lack of sufficient data or appropriate approaches to detect real patterns. If our sample size had been much larger, if we had used another technique for integrating variables (or just the right tweak on our methods), or if we had better biomarkers, we might well have detected a clear pattern. But if this is the case, it is not encouraging for our ability to detect such signals in future studies: as can be seen here, the more methods are tried, the harder it becomes to correctly identify a true positive among the false positives certain to arise through multiple testing. If the detection of a true signal depends precisely on details such as the choice of biomarkers or how DM is centered, and if these details vary (as is likely) across species, populations, and conditions, future results, both positive and negative, must be viewed with a healthy grain of salt.

Even if our negative results are due to the lack of appropriate methods, they still have important biological implications: if there were a clear, simple, straightforward signal, we would have detected it. Our birds showed a wide range of reproductive success and reproductive effort: some individuals made only one breeding attempt, had no breeding productivity, and did not return the following year, whereas other individuals reared up to 10 chicks from two broods, returned the following year, and reared up to 8 more chicks from two further broods. This implies ample opportunity to detect associations. Many of our biomarkers have shown associations with aspects of life histories or performance in other studies [36-41], and, as much as possible given physiological complexity, there was solid physiological theory behind their selection. The clear implication is that associations between physiology and “quality”-related measures are either highly variable or largely absent, both in our study and more generally. An increasing number of studies highlight the complexity of the physiology underlying the biomarkers measured in ecophysiology studies. For example, correlation structures among immune markers vary even within populations over time and across habitats, as well as across species and at different taxonomic levels [12, 13, 23]. Such complexity is to be expected: most of the markers used play multiple physiological roles, and changes in their levels could be due to many environmental or physiological factors, such that any observed associations result not from meaningful causality but from complex contingency.

In this context, it is unsurprising that the univariate methods in the companion paper produced little quality signal despite some evidence for systematic annual- and breeding-stage-dependent variation in physiological traits, but more surprising that the multivariate methods here did not. Indeed, other research on DM has shown it to be broadly robust to choice of component biomarkers as long as 10+ are included [25], and this was confirmed here. Various versions of DM used here were highly correlated, suggesting that the details of calculation method were not the limiting factor. Rather, we see our findings as a true absence of strong association between “dysregulation” (as measured by DM) and reproductive variables in this free-living population. In our original paper proposing DM as an ecophysiology measure, we found clear associations between DM and both metabolic capacity and a foot inflammation score, each in the expected direction [27]. However, those performance variables were more directly related to health status, not fitness.

While it is certainly possible that, despite our previous finding with the red knots, DM is not capturing a signal relevant to anything important in the starlings, another intriguing possibility is that DM is accurately capturing health status or condition, but that condition has no straightforward association with the reproductive variables we measured here. This might be due to non-monotonic relationships or alternative strategies. For example, perhaps older individuals or those in failing health make a large terminal investment in reproduction [42, 43], obscuring a more general positive association between condition and reproductive success. Alternatively, perhaps highly dysregulated individuals choose not to reproduce and thus were excluded from our study. If this is the case, it would suggest that DM specifically, and measures of condition more generally, may not show interesting relationships with reproductive success: the individuals in the worst condition do not breed, and the residual variation in condition among breeders may be small. It might also be confounded by a hard-to-disentangle negative feedback loop: a positive effect of physiological state on effort, a positive effect of effort on physiological costs, and a negative effect of physiological costs on physiological state.

One potential reason for the discrepancy between this study and our previous work on DM is the substantial expected and observed physiological variation within our population. While our total sample was relatively large, sub-samples by year and breeding stage were not, and many of our measures differed markedly across these sub-groups. In contrast, red knots were captive during that study and were measured ∼12 times each, reducing variation. Likewise, the human cohorts (mostly elderly) that we have used [25, 28, 29, 31, 32] live in stable environments in developed countries and are not undergoing seasonal breeding cycles. The principle of DM relative to homeostatic control requires a relatively homogeneous population, and this was likely not the case here. Our best guess is thus that DM might still perform well for physiologically homogeneous populations (same breeding stage, parasite pressure, etc.) but may break down as variation in optimal physiological state increases, perhaps related to more highly variable ecological context (as well as life history context).

We are aware that some readers may find our analyses either intimidating or undirected and too exploratory. If we had claimed to have a clear positive result, it would indeed be critical to consider the possibility of a false positive in relation to multiple testing. However, we also argue that current knowledge about the relationship between physiology and fitness is insufficient to provide clear testable hypotheses, necessitating some level of exploratory analysis. In this sense, the core take-home message is that even with such an exploratory approach and so much effort put into the statistical analyses, there is no clear biological signal. This is not an encouraging message for ecophysiology, but it is not one that can be ignored simply because it is inconvenient. Going forward, it will be important to try to replicate our negative finding using other integration approaches, species, and biomarkers. There may be particular factors related to our study that caused a negative result for an approach that might be promising more broadly. The positive result in red knots was highly robust, not an artefact of data massaging. Nonetheless, we feel this study provides a strong warning to eco-physiologists hoping to use physiological measures to quantify body condition or individual quality, particularly in cases where there may be substantial physiological heterogeneity in the sample.

## Acknowledgments

Thanks to Sarah Gray and James Hou for help with wing measurements and reticulocyte counts. Kevin Matson, Chris Harris, Oliver Love, David Costantini, Sarah Guindre-Parker, Christopher Guglielmo and David Swanson all provided invaluable advice regarding troubleshooting assays. Special thanks to Allison Cornell for invaluable help with field and lab work.

---

**S1 Appendix.** Supplemental methods, tables and figures.

## Abbreviations

AG: Agglutination score from Nab assays.
AG-lysis PCA: PCA on AG and lysis variables, reflecting the immune function.
BSF: Brood size at fledging.
BSF1: BSF for first brood only.
BSF2: BSF for second brood only.
BSF sum 2yrs: Total number of offspring that fledged in both years.
CK: Creatine kinase concentrations, (U/L).
Cort: Corticosterone (ng/mL).
DM: A measure of body condition based on statistical distance (Mahalanobis distance).
DM V0: DM calculated using the mean as the centroid for each biomarker.
DM V1-V4: DM versions based on a priori hypotheses, either using the mean + 3 standard deviations (high) or the mean – 3 standard deviations (low).
DM V5: DM version generated by random choice of mean, high (mean + 3 standard deviations), or low (mean - 3 standard deviations) for the centroid definition.
dROMs: Reactive oxygen metabolites (mg H2O2/dL).
NEFA: Non-esterified fatty acids (mmol/L).
OXY: Antioxidants titers (µmol HClO/mL).
OXY-dROMs PCA: PCA on antioxidants (OXY) and oxidative stress measures (dROMs).
PCA: Principal components analysis.
PCA1: First PCA axis.
PCA2: Second PCA axis.
PCA3: Third PCA axis.
PCA aerobic capacity: PCA on physiological variables representing aerobic and metabolic capacity (hematocrit, hemoglobin, reticulocytes, and corticosterone).
PCA deviational: PCA on all physiological variables, centered and reduced by year and stage, using the logarithm of their absolute values.
PCA directional: PCA on all physiological variables, centered and reduced by year and stage.
PCA metabolism/energy: PCA on physiological variables representing intermediary metabolism and energy supply (NEFA, triglycerides, uric acid, and glucose).
PCA oxidative stress: PCA on physiological variables representing oxidative stress and muscle damage (CK, OXY, and dROMs).
Reproduction PCA: PCA including mean fledge mass, brood size at day 6, BSF, and BSF sum per year.

